# FrameDiPT: SE(3) Diffusion Model for Protein Structure Inpainting

**DOI:** 10.1101/2023.11.21.568057

**Authors:** Cheng Zhang, Adam Leach, Thomas Makkink, Miguel Arbesú, Ibtissem Kadri, Daniel Luo, Liron Mizrahi, Sabrine Krichen, Maren Lang, Andrey Tovchigrechko, Nicolas Lopez Carranza, Uğur Şahin, Karim Beguir, Michael Rooney, Yunguan Fu

## Abstract

Protein structure prediction field has been revolutionised by deep learning with protein folding models such as AlphaFold 2 and ESMFold. These models enable rapid *in silico* prediction and have been integrated into *de novo* protein design and protein-protein interaction (PPI) prediction. However, biologically relevant features dependent on conformational distributions cannot be estimated with these models. Diffusion models, a novel class of generative models, have been developed to learn conformational distributions and applied to *de novo* protein design. Limited work has been done on protein structure inpainting, where a masked section is recovered by simultaneously conditioning on its sequence and the rest of the structure. In this work, we propose **FrameD**iff **i**n**P**ain**T**ing (FrameDiPT), a generalised model for protein inpainting. This is important for T-cells given the hyper-variability of the complementarity determining region (CDR) loops. We evaluated the model on CDR loop design for T-cell receptors and achieved comparable prediction accuracy to ProteinGenerator and RFdiffusion with limited training data and learnable parameters. Different from deterministic structure prediction models, FrameDiPT captures the conformational distribution at different regions and binding states, highlighting a key advantage of generative models. The model and inference code have been released^1^.

## 1 Introduction

Proteins play an essential part in almost all cellular processes. Accurate modelling of protein structure is important to assess the behaviour of existing and *de novo* proteins. While models such as AlphaFold 2 [Jumper et al., 2021, Evans et al., 2021], RoseTTAFold [Baek et al., 2021] and ESMFold [Lin et al., 2022] have revolutionised computational protein modelling with high-quality predictions, their deterministic nature does not reflect the dynamic nature of proteins. Diffusion denoising models [Sohl-Dickstein et al., 2015, Ho et al., 2020], a novel class of generative models, have achieved superior performance in image synthesis. RFdiffusion [Watson et al., 2023] integrated the diffusion model with a pre-trained RoseTTAFold to perform *de novo* protein design, motif scaffolding and binder design. While recent works have leveraged diffusion models for conformational distribution tasks by training on molecular force fields [Abdin and Kim, 2023, Zheng et al., 2023], limited work has been done on protein structure inpainting tasks where only a subset of the residues are of interest. By fixing the majority of residue positions, we sample the conformational distribution in the area of interest while avoiding incorrect global structure predictions.

In this work, we focus on T-cell receptors (TCRs) and peptide-major histocompatibility complexes (pMHCs), which are crucial for cell-mediated immunity. The complementarity determining region (CDR) loops of TCRs, especially the CDR3 loops, are highly variable and thereby able to bind with different pMHCs [Minguet et al., 2007, Xu et al., 2020]. Understanding the conformational distribution is therefore beneficial to downstream tasks such as TCR maturation and binding prediction. We choose to model the distribution of CDR3 loops and keep the rest of the structure fixed as contextual information. This differs from DiffAB [Luo et al., 2022] which designs antibody CDR sequences and structures given antibody-antigen frameworks, since, in our inpainting task, the amino acid sequences are known. We extended FrameDiff [Yim et al., 2023], an SE(3) diffusion model for *de novo* protein backbone generation, to generic protein structure inpainting, a model we term **FrameD**iff **i**n**P**ain**T**ing (FrameDiPT). After training for 2 GPU-weeks on 32K monomer structures, an 18M parameter FrameDiPT model reached satisfying prediction accuracy compared to deterministic models including AlphaFold 2, ESMFold as well as diffusion-based models such as RFdiffusion and ProteinGenerator [Lisanza et al., 2023]. Importantly, the compared methods were trained on larger datasets including TCR-like structures, while FrameDiPT training intentionally excluded all structures similar to TCRs and antibodies. FrameDiPT also captured conformational distribution differences at different regions and binding states. The model and inference code have been released^1^.

## 2 Related work

Diffusion models [Sohl-Dickstein et al., 2015, Ho et al., 2020] were first applied to image generation, where images are synthesised via progressive denoising. Different from images which have no limitation in sampling space, molecules and proteins have intrinsic geometric constraints on bond angles and lengths. AlphaFold 2 [Jumper et al., 2021] represents each residue as a rigid frame thus preventing structural violations. Similarly, heavy atom side chain positions can be described by dihedral angles in lieu of bond lengths [Baek et al., 2021]. With these rigid representations, diffusion models on the Riemannian manifolds SO(2) and SO(3) [Huang et al., 2022, Leach et al., 2022] are applied to proteins to ensure plausible structure sampling.

While small molecule designs [Shi et al., 2021] model all atoms, most protein diffusion models focus on *de novo* protein design where only backbone atoms [Watson et al., 2023, Yim et al., 2023, Wu et al., 2022] are generated, reducing the model complexity and computational costs. Full-atom structures can be obtained by sampling a sequence via inverse folding models such as ProteinMPNN [Dauparas et al., 2022], and then predicting structure using protein folding models. Alternatively, full-atom generation models such as cg2all [Heo and Feig, 2023] can be used to generate full structures based on coarse-grain representations. Other works such as Chroma [Ingraham et al., 2022], which attempts to preserve the average bond angle and radius of gyration through an anisotropic diffusion process, and EigenFold [Jing et al., 2023] which performs diffusion on a harmonic decomposition of the protein chain instead of directly in the coordinate space of residues, attempt to integrate physical priors into the diffusion processes. Some work also attempts to learn sequence-structure joint distributions and thus performs protein sequence-structure co-design [Zhang et al., 2023, Anand and Achim, 2022, Luo et al., 2022, Chu et al., 2023]. However, there is limited research focusing on the task of protein structure inpainting. Wang et al. [2022] proposed a deep learning model for scaffolding protein functional sites, however, with a deterministic model. RFdiffusion targets various tasks including motif scaffolding but relies on ProteinMPNN to generate sequence. Similarly, Gao et al. [2023] proposed a language diffusion model DiffSDS for unknown-sequence protein backbone inpainting.

## 3 Method

### 3.1 FrameDiff

Yim et al. [2023] introduced FrameDiff, a graph-based SE(3)-equivariant neural network, which achieves comparable performance to RFdiffusion for *de novo* protein backbone design with much less training data and without a pre-trained structure prediction network. FrameDiff uses the backbone rigid-frame representation T with an additional torsion angle *ψ* to determine the position of the backbone carbonyl oxygen atom. The forward diffusion process is performed on the rigid-frame representation, i.e. the translation **X** given by the coordinates of the *C*_*α*_ atom and the rotation **R** defined by the frame formed by the *N* − *C*_*α*_ − *C* atoms for each residue. The torsion angle *ψ* is not involved in the diffusion process but is predicted by the model. The backward diffusion process is defined by denoising score matching [Vincent, 2011]. Training losses consist of both the rigid-frame losses and the atom-level losses. We refer the readers to Yim et al. [2023] for further details.

### 3.2 FrameDiPT

We extend FrameDiff to **FrameD**iff **i**n**P**ain**T**ing (FrameDiPT) for protein structure inpainting with the following modifications.

#### Randomly mask a contiguous region for diffusion

For all training monomers, a contiguous region of 8-50 amino acids is randomly selected and the diffusion process is applied only in this region. For the input node features, the diffusion timestep `t` is set to 10^−5^ in the fixed region to indicate the residues are not being diffused. For convenience, we call the randomly selected contiguous region the “diffused region” and the remaining part of the structure the “context region”.

#### Add amino acid types as node features

In contrast to *de novo* protein design, the amino acid sequence is given in the inpainting task. An extra node feature `aatype` is concatenated to the original node features of FrameDiff. For a protein with *N*_*res*_ residues, the `aatype` node feature is of shape (*N*_*res*_, 21) containing 20 standard amino acid types and 1 for unknown amino acid type. Similarly, edge features are modified accordingly.

#### Training loop

During the training loop, the diffusion noising process is only performed in the diffused region. The resulting noised structure is then given as input to the model to predict the original structure. Loss is only computed over the diffused region. Different from FrameDiff, we train the model on clustered data with a sequence similarity threshold of 90%. In each epoch, only one structure is randomly sampled from each cluster. This approach outperforms FrameDiPT trained with non-clustered data (Appendix B).

#### Inference loop

In the inference loop, the context region along with the sampled random noise in the diffused region are given as the initial state. In each inference step, the model predicts the backbone rigid frames 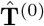from **T**^(*t*)^, where **T**^(*t*)^ represents the rigid frames at time *t* and *t* = 0 corresponds to the ground truth. A reverse diffusion step is performed to get **T**^(*t*−*dt*)^ from **T**^(*t*)^ and 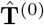 where *dt* is the step size. Then the context region in **T**^(*t*−*dt*)^ is replaced by the original structure, to ensure the context region stays fixed. Finally, we extended our model to run inference on multimers by adding a residue gap of 200 between different chains, following the trick used in Motmaen et al. [2023].

## 4 Experiment setting

### Training

We train FrameDiPT using data from RCSB Protein Data Bank (RCSB PDB)^2^ (Appendix A) with the same hyperparameters as FrameDiff. It was trained for 2 weeks with 1 NVIDIA A100 GPU for 95 epochs with a length batching strategy. Each batch contains a collection of different diffused instances of the same backbone structure, and the number of samples per batch adapts to the sequence length of the structure with a maximum batch size of 128.

### Evaluation

A TCR and TCR:pMHC dataset has been curated for evaluation (Appendix C). It contains three splits: 21 unbound TCR structures; 62 TCR:pMHC class I complexes; and 18 TCR:pMHC class II complexes. For each TCR or TCR:pMHC sample, the CDR3 loop in both TCR alpha and beta chains are masked as the diffused region and 100 inference steps are performed to get the final prediction. `cg2all`^3^ is used to convert the predicted structure to an all-atom structure. We report the root-mean-square-deviation (RMSD) on the backbone and full-atom structure for evaluation. For diffusion models, we developed a sample selection strategy using kernel density estimation. A Gaussian kernel with standard deviation of 30Å is fitted over the carbon alpha coordinates of the inpainted region to estimate density. The sample corresponding to the highest density is used as a proxy for the “most-likely” sample.

### Baseline diffusion models

ProteinGenerator performs diffusion in sequence space and structure is predicted via RoseTTAFold given the generated sequence. We adapted ProteinGenerator to the inpainting task by supplying the input sequence. RFdiffusion is used by masking out both the structure and sequence of TCR CDR3 loops, thus full-atom RMSD is not applicable. Empirically, we found providing sequences to RFdiffusion deteriorated the performance since the model was not trained in this mode.

## 5 Results and discussions

### Comparison to protein diffusion models

We compared FrameDiPT with existing diffusion models, ProteinGenerator and RFdiffusion, for inpainting tasks (Table 1). The original FrameDiff is not applicable as it does not take any sequential or structural information as input. Across five samples, FrameDiPT achieved median RMSD of 2.70Å, 2.18Å, and 2.91Å on unbound TCR, bound TCR with pMHC-I, and pMHC-II, respectively. This performance is comparable with ProteinGenerator and RFdiffusion which have larger networks and have been trained on larger datasets (Appendix D). A signed Wilcoxon paired two-sided rank test was performed which indicated no statistically significant difference between results. Importantly, the training of FrameDiPT explicitly excluded TCR:pMHC and antibody structures (Appendix A), demonstrating the strong generalization capacity of the proposed method. The RMSD is further reduced across all datasets when generating 25 samples (Table 5). This indicates that FrameDiPT is capable of sampling structures that are specified in crystal structures. It is important to highlight that, although the structures were evaluated using RMSD, RMSD to X-ray structure is not the perfect metric as the crystalised conformation represents only a snapshot of the dynamic.

**Table 1:** Backbone and full-atom RMSD comparison. A signed Wilcoxon paired two-sided rank statistical test between FrameDiPT and ProteinGenerator is performed at significance level p-value < 0.05. Underline means significantly better than ProteinGenerator which outperforms RFdiffusion.

| Method | RMSD | TCR | TCR:pMHC-I | TCR:pMHC-II |
| --- | --- | --- | --- | --- |
| ProteinGenerator | Backbone | $2.87 \pm 0.48$ | $2.34 \pm 0.73$ | $3.01 \pm 0.55$ |
| | Full-atom | $3.58 \pm 0.62$ | $3.24 \pm 0.79$ | **$3.59 \pm 0.39$** |
| RFdiffusion | Backbone | $3.23 \pm 0.93$ | $3.32 \pm 0.72$ | $3.81 \pm 0.91$ |
| FrameDiPT | Backbone | **$2.70 \pm 0.43$** | **$2.18 \pm 0.45$** | **$2.91 \pm 0.54$** |
| | Full-atom | **$3.48 \pm 0.52$** | **$2.93 \pm 0.59$** | $3.90 \pm 0.46$ |

### Capturing conformational distributions

While CDR3 loops are highly flexible, the N- and C-terminal flank regions should be more constrained, as they are mainly beta strands. We thus performed inpainting on these flanks of the same length as the CDR3 loop to compare the variance of generated samples, which is defined as the average inter-sample backbone RMSD (Table 2). Lower backbone RMSD and sample variance were observed in the flanks. Notably, the C-terminal flank has a smaller variance than N-terminal, correctly reflecting how the C-terminus fully encompasses a beta strand while the N-terminal flank starts on the small loop leading the beta strand before the CDR3 loop. The correlation between normalised carbon-alpha B-factors and sampling variances (Figure 9b) is 0.366. The moderate correlation is not surprising, as B-factors aggregate different effects from experimental uncertainty to structural flexibility in a variable manner between structures.

**Table 2:** Backbone RMSD and sample variance of different diffused regions. N- and C-terminal flanks have lower RMSD and variance, consistent with structural properties of the diffused regions.

| Diffused region | Metric | TCR | TCR:pMHC-I | TCR:pMHC-II |
| --- | --- | --- | --- | --- |
| CDR3 | Backbone RMSD | $2.70 \pm 0.43$ | $2.18 \pm 0.45$ | $2.91 \pm 0.54$ |
| | Variance | $1.60 \pm 0.20$ | $1.58 \pm 0.34$ | $1.85 \pm 0.34$ |
| N-terminal flank | Backbone RMSD | $0.89 \pm 0.23$ | $1.59 \pm 0.48$ | $1.71 \pm 0.39$ |
| | Variance | $0.92 \pm 0.37$ | $0.96 \pm 0.23$ | $0.93 \pm 0.30$ |
| C-terminal flank | Backbone RMSD | $0.71 \pm 0.12$ | $0.69 \pm 0.08$ | $0.76 \pm 0.14$ |
| | Variance | $0.26 \pm 0.10$ | $0.32 \pm 0.13$ | $0.38 \pm 0.10$ |

### Conformation changes upon binding

The CDR conformations differ in the bound and unbound states [Armstrong et al., 2008]. This is due to intermolecular forces and steric constraints induced by the pMHC complex influencing the position of loop residues. Pairs of structures (6 for pMHC-I and 2 for pMHC-II) are found that represent unbound and bound states of the same TCRs. For each structure, 100 samples were generated, with inpainting applied to the CDR3 loops on the alpha and beta chains. Figure 1 shows a clear separation between unbound and bound samples in terms of CDR3 loop conformations for 2BNU and 2BNQ, with significant differences between distribution centroids. More examples can be found in Appendix E.3. This underlines the strength of diffusion models in capturing different conformations, which could be useful for downstream binding classification tasks.

**Figure 1.**
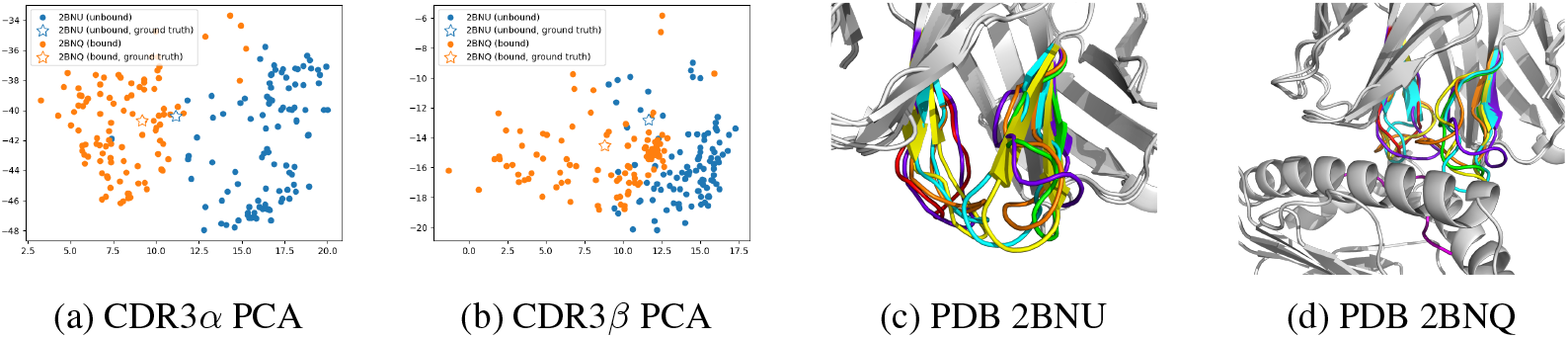
Plots of the first two principal components of PCA describing a) CDR3*α* and b) CDR3*β C*_*α*_ position loop conformations with ground truth marked as a star. A clear distinction between samples drawn from the unbound (2BNU) and bound (2BNQ) conformations can be seen on the alpha chain. A clear difference in modes can be seen on the beta chain. c) 2BNU and d) 2BNQ visualizations with context structure, ground truth CDR3 alpha, ground truth CDR3 beta and peptide; FrameDiPT predictions in cyan, yellow and orange alongside ESMFold prediction.

For example, sampling loop conformations of different or mutated bound TCR:pMHC complexes and evaluating them through energy-scoring methods could help to discriminate weak and strong binders.

### Quantifying uncertainty

Figure 9a shows the correlation between the median backbone RMSD and the variance of generated samples. Interestingly, when the sampled structures are different between them, the samples themselves may be inaccurate. The variance can thus be used as a metric to analyse the overall quality of generated samples for a test instance.

### Comparison to deterministic protein folding models

FrameDiPT has been compared with pre-trained deterministic protein folding models such as AlphaFold 2 (Table 7). AlphaFold 2 with custom templates where CDR3 loops were masked had lower RMSD to the ground truth crystal structures. The difference to FrameDiPT is significant, indicating further room for improvement.

## 6 Conclusion

In this work, we proposed a novel inpainting task for protein structure generation. We trained the proposed FrameDiff inpainting (FrameDiPT) model on 32K monomer structures and evaluated it on TCR CDR3 loop design. With no TCR and antibody structures present in the training data and only 18M parameters, FrameDiPT achieved similar RMSD to other TCR-aware pre-trained large diffusion models such as ProteinGenerator and RFdiffusion. Despite only being trained on monomers, FrameDiPT could capture the conformational distribution of the diffused region and the TCR:pMHC binding interaction. While FrameDiPT is able to sample structures close to the crystal structures, the RMSD remains significantly higher than AlphaFold 2 which has a larger network and larger training set. In the future, FrameDiPT could be improved with more training data and scaling up the network to close the gap. Moreover, downstream applications such as TCR:pMHC binding classification can be considered.

## A Training data

Data used to train FrameDiPT model was downloaded from RCSB PDB with the following data cleaning procedure. Only X-ray structures were kept, i.e. the structures from non-X-ray assays, or ModelArchive^4^ and the AlphaFold 2 predicted structures were removed. The training data cleaning process also included the removal of structures belonging to any of the following categories: 1) have only non-standard residues; 2) have a resolution larger than 9Å; 3) have a single amino acid accounting for more than 80% of the structure; 4) have more than 4950 residues. In the end, any structures that could not be parsed by `biopython`^5^ were removed.

The data cleaning process involved also removing samples that are similar to the TCR test data to avoid data leakage. Similar samples to TCR test data are identified using the 70% sequence similarity clusters. A cluster is considered as “leaking” if it contains any chain from the TCR test data. Afterwards, all samples from “leaking” clusters are removed from the training set. Such removal ensures that structures in the training set have a maximum sequence similarity of 70% compared to any structure in the TCR test data.

We followed the same data processing procedure as in the original FrameDiff, which leads to 32K monomers for training and the training strategy on clustered data leads to 9K clusters for the 32K monomers.

## B Training strategy

The training on clustered monomers with 9K clusters is evaluated against baseline training on all 32K monomers for both *de novo* protein design model FrameDiff and inpainting model FrameDiPT. Figure 2 shows self-consistency RMSD, which is the RMSD between the generated backbone and ESMFold predictions of the ProteinMPNN generated sequences, for different designed lengths. The model trained on clustered data shows consistently better results than the baseline model. Table 3 shows median backbone RMSD on CDR3 loop design where better performance is observed with training on clustered data.

**Table 3:** Backbone RMSD comparison between baseline training and training on clustered data. A signed Wilcoxon paired two-sided rank statistical test between baseline and clustered training is performed at significance level p-value < 0.05. Underline means significantly better.

|  | Baseline | Clustered |
| --- | --- | --- |
| TCR | $2.77 \pm 0.47$ | **<u>$2.70 \pm 0.43$</u>** |
| TCR:pMHC-I | $2.86 \pm 0.74$ | **<u>$2.18 \pm 0.45$</u>** |
| TCR:pMHC-II | $3.15 \pm 0.46$ | **<u>$2.91 \pm 0.54$</u>** |

**Figure 2.**
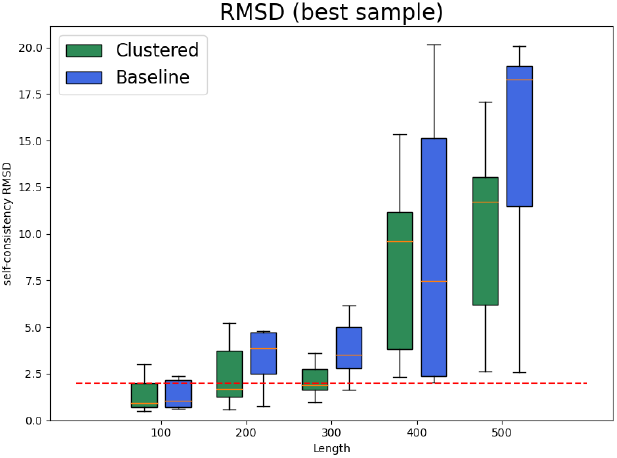
Best sample self-consistency RMSD of *de novo* protein design with the baseline model and the model trained on clustered data. The latter shows consistently better performance for all designed lengths.

## C Evaluation data

Curated sets of high-resolution, annotated structures of TCRs and TCR:pMHC complexes without any other companion proteins were assembled and fetched from the RCSB. First, the lists of PDB IDs for the different dataset types were fetched from the Structural T-Cell Receptor Database (STCRDab) [Leem et al., 2017]. The corresponding structures were downloaded from the RCSB and only X-ray structures with resolution < 3.5Å were kept. Then, the PDBe REST API was used to map the structures’ chains to UniProt IDs, when available. The UniProt metadata was used to label the chains in each structure (e.g. TCR alpha or beta chain, peptide, MHC alpha or beta chain) based on keyword and gene name matching. Other proteins or not annotated ones were flagged as such. Only structures with the expected TCR, peptide, or MHC chains^6^ were kept; structures containing other proteins, or unlabelled ones were filtered out.

## D Experiment settings

Experiment settings including number of model parameters, training time and training data of different models are summarised in Table 4. As ProteinGenerator and RFdiffusion are fine-tuned RoseTTAFold model, the experiment setting for RoseTTAFold is reported. FrameDiPT model has significantly fewer parameters and is trained with much less data and training time.

**Table 4:**
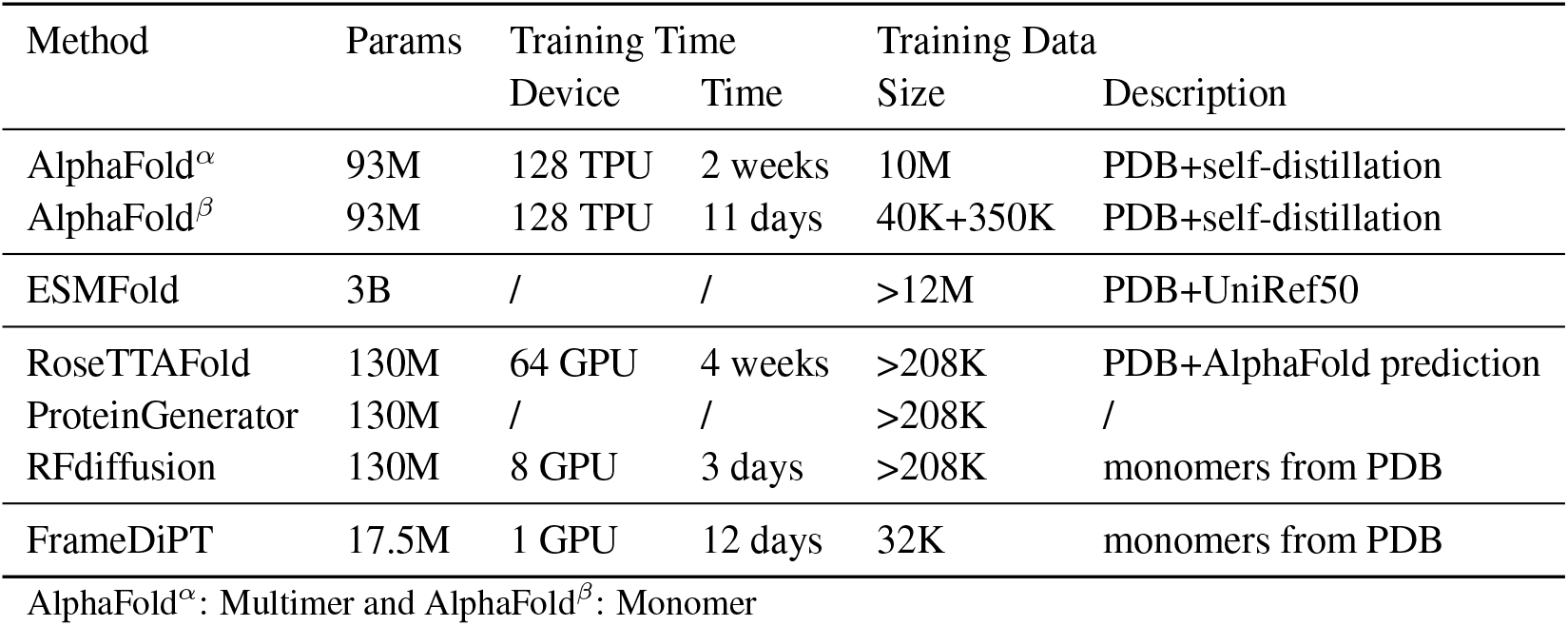
Experiment setting comparison.

**Table 5:**
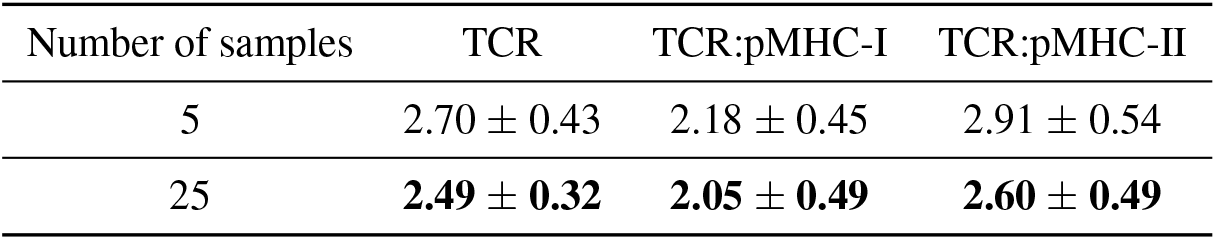
Backbone RMSD w.r.t. number of generated samples.

| Number of samples | TCR | TCR:pMHC-I | TCR:pMHC-II |
| --- | --- | --- | --- |
| 5 | 2.70 ± 0.43 | 2.18 ± 0.45 | 2.91 ± 0.54 |
| 25 | **2.49 ± 0.32** | **2.05 ± 0.49** | **2.60 ± 0.49** |

## E Further results and discussions

### E.1 Generating specific conformation

### E.2 Capturing conformational distributions

Backbone RMSD per residue is also computed to evaluate how our model performs at different residue positions and we observe different patterns for different diffused regions. Figure 4 visualises an example (PDB 1KGC) of generated samples for CDR3 N-terminal and C-terminal flanks, which shows consistent structural properties. Figure 3 shows backbone RMSD per residue of CDR3, N-terminal and C-terminal flanks of CDR3. The CDR3 loop shows larger RMSDs in the middle of the loop. For N-terminal flank, the positions close to CDR3 loop usually consist of beta strands for which small RMSDs are obtained while for the positions further away from CDR3 loop, more potential conformations are predicted therefore leading to higher RMSD. For the C-terminal flank, the RMSD is consistently low which is coincident with end of the loop becoming a beta strand. Position 3 is an exception showing a local increase of RMSD, consistent with a kink following the loop at the start of the beta strand.

**Figure 3.**
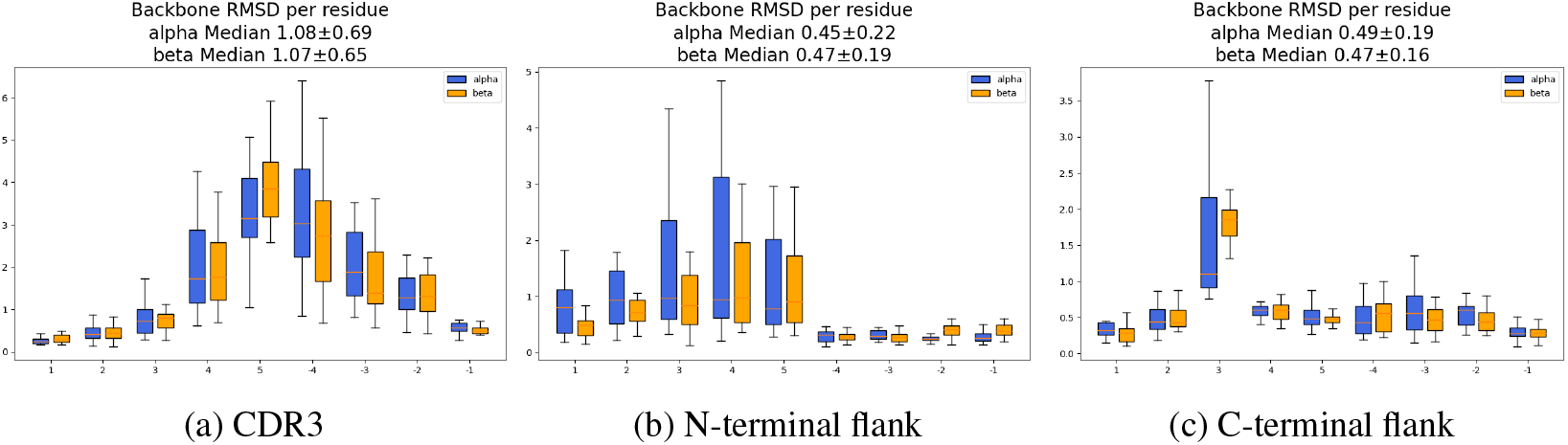
Backbone RMSD per residue on TCR dataset of (a) CDR3, (b) N-terminal flank and (c) C-terminal flank. CDR3 loop shows greater RMSD in the middle of the loop while N-terminal flank shows smaller RMSD at positions close to CDR3 and C-terminal flank shows small RMSD in general except the third position.

**Figure 4.**
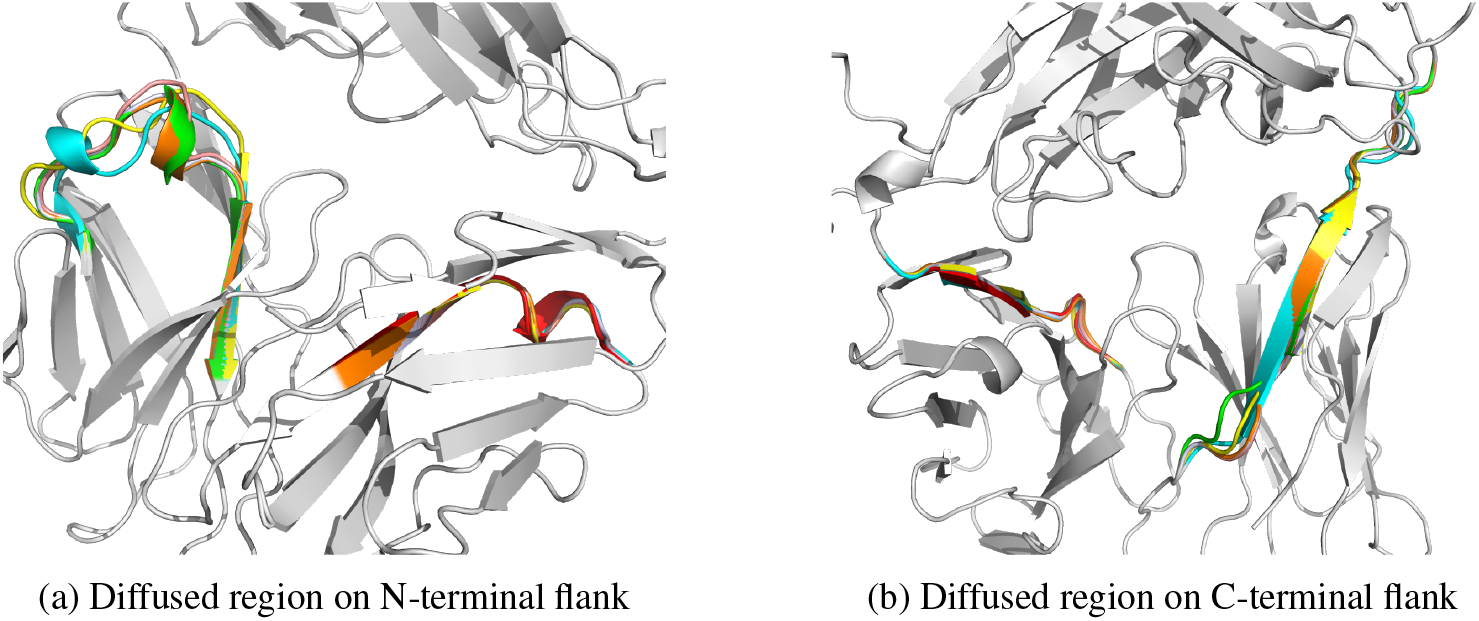
PDB 1KGC with context structure, ground truth alpha, ground truth beta and FrameDiPT predictions in other colors for (a) diffused region on N-terminal flank and (b) diffused region on C-terminal flank. Structural properties correspond to backbone RMSD per residue shown in Figure 3 where positions with smaller RMSD are usually beta strands and those with bigger RMSD are usually loops, especially the 3rd position of C-terminal flank corresponds to a kink in the loop structure.

**All CDR loops diffusion** Loops are usually flexible structures while different loops could have different structure flexibility, for example CDR3 loops are the most variable w.r.t. CDR1 and CDR2 loops in TCR chains. We also performed diffusion on all the three CDR loops and compared the backbone RMSD and sample variance in Table 6. Smaller backbone RMSD and sample variance are obtained for CDR1 and CDR2 loops.

**Table 6:**
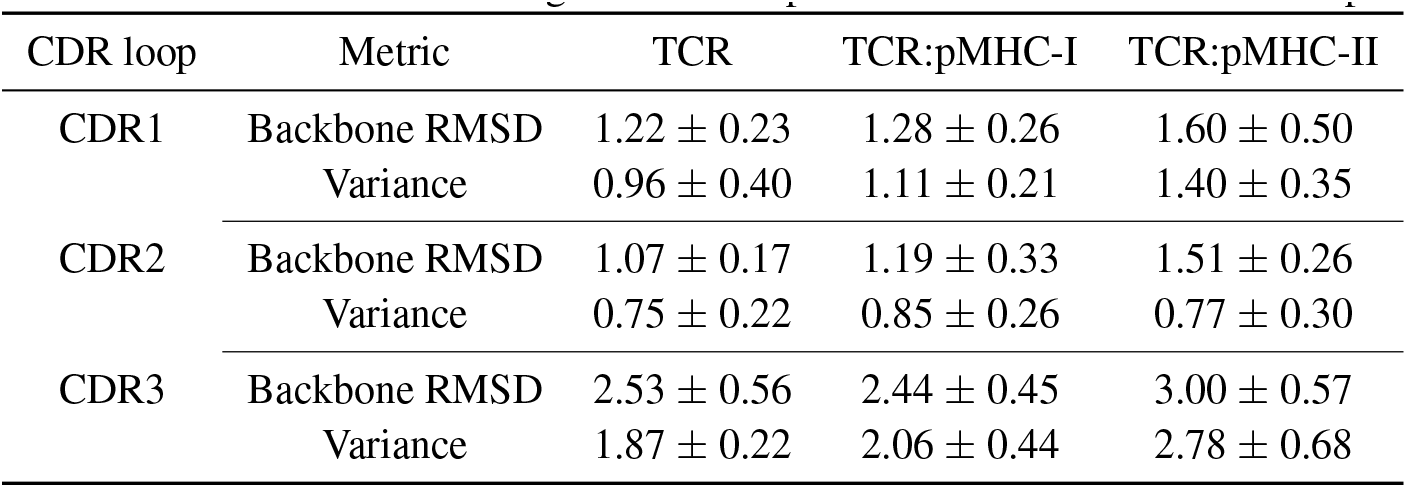
Backbone RMSD and generated sample variance of different CDR loops.

**Table 7:** Backbone and full-atom RMSD comparison of TCR CDR3 loop design. A signed Wilcoxon paired two-sided rank statistical test between FrameDiPT and the best AlphaFold model is performed at significance level p-value < 0.05. Underline means significantly different from the best AlphaFold model.

| Method | RMSD | TCR | TCR:pMHC-I | TCR:pMHC-II |
| --- | --- | --- | --- | --- |
| AlphaFold<sup>α</sup> | Backbone | $2.60 \pm 0.68$ | $2.46 \pm 0.68$ | $2.33 \pm 0.65$ |
| | Full-atom | $3.15 \pm 0.54$ | $2.97 \pm 0.76$ | $2.91 \pm 0.65$ |
| AlphaFold<sup>β</sup> | Backbone | **$1.75 \pm 0.68$** | $1.58 \pm 0.51$ | $1.75 \pm 1.08$ |
|  | Full-atom | **$2.20 \pm 0.72$** | **$2.27 \pm 0.75$** | **$2.04 \pm 1.03$** |
| AlphaFold<sup>γ</sup> | Backbone | $2.03 \pm 0.68$ | **$1.34 \pm 0.48$** | **$1.37 \pm 0.43$** |
| | Full-atom | $2.49 \pm 0.66$ | **$2.27 \pm 0.87$** | $2.10 \pm 0.65$ |
| ESMFold | Backbone | $2.58 \pm 0.86$ | $2.13 \pm 0.42$ | $2.27 \pm 0.85$ |
| | Full-atom | $3.24 \pm 0.80$ | $2.83 \pm 0.66$ | $2.41 \pm 0.70$ |
| FrameDiPT (25 samples) | Backbone | <u>$2.49 \pm 0.32$</u> | <u>$2.05 \pm 0.49$</u> | <u>$2.60 \pm 0.49$</u> |
|  | Full-atom | <u>$3.45 \pm 0.47$</u> | <u>$2.83 \pm 0.65$</u> | <u>$3.26 \pm 0.56$</u> |

AlphaFold <sup>$\alpha$</sup> : AlphaFold Multimer with searched templates from PDB70.
AlphaFold <sup>$\beta$</sup> : AlphaFold Multimer with custom templates by masking CDR3 loops.
AlphaFold <sup>$\gamma$</sup> : AlphaFold Monomer with custom templates by masking CDR3 loops.

### E.3 Conformation change upon binding

**Figure 5.**
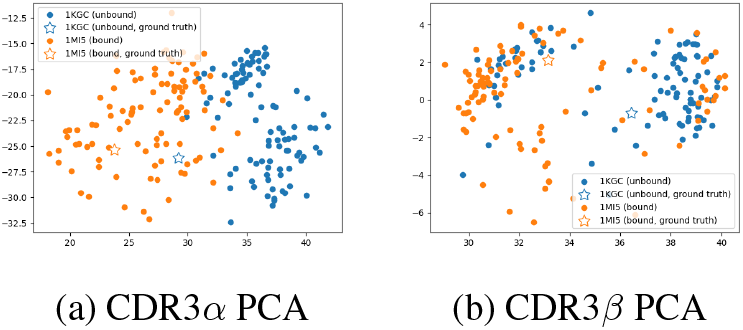
Conformational distributions of 1KGC (unbound) and 1MI5 (bound)

**Figure 6.**
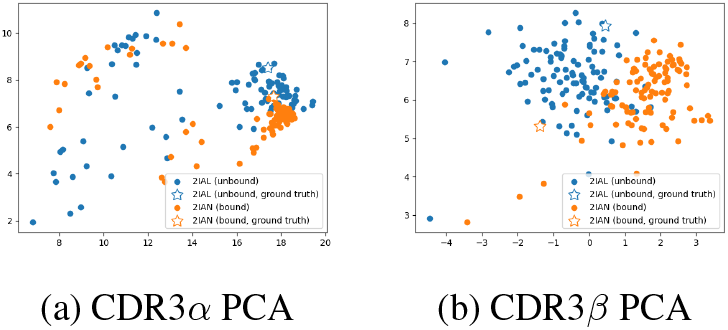
Conformational distributions of 2IAL (unbound) and 2IAN (bound)

**Figure 7.**
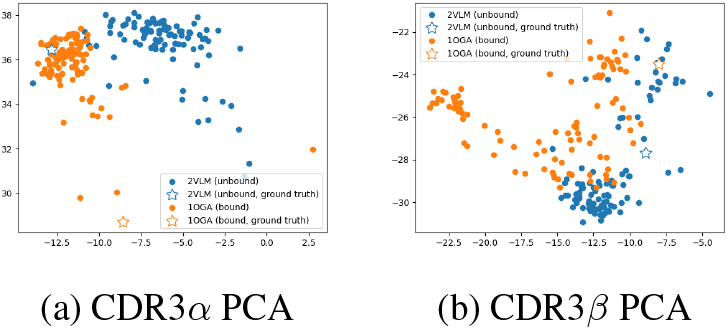
Conformational distributions of 2VLM (unbound) and 1OGA (bound)

**Figure 8.**
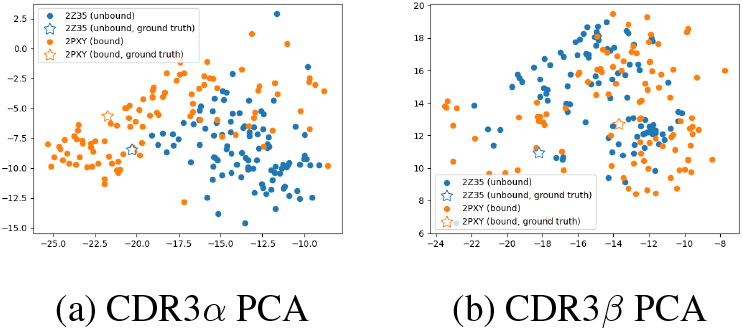
Conformational distributions of 2Z35 (unbound) and 2PXY (bound)

### E.4 Quantifying uncertainty

We analysed the correlation between backbone RMSD and sampling variance (Figure 9a) and between normalised carbon-alpha B-factors and sampling variance (Figure 9b). Though we performed a standard normalisation of B-factors over the whole protein structure to remove intrinsic factors, no evident correlation between B-factors and sampling variance was observed.

### E.5 Comparison to deterministic protein folding models

**Figure 9.**
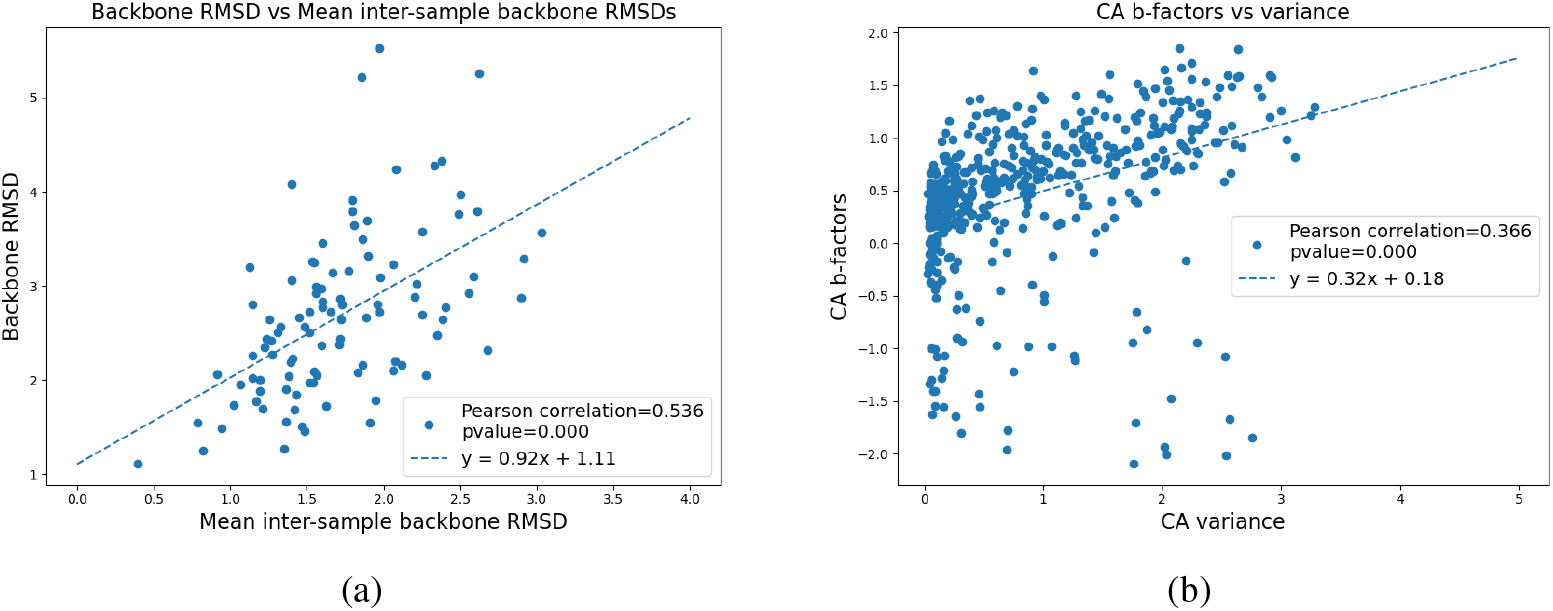
Correlation between a) median backbone RMSD and the variance of generated sample which is computed as mean inter-sample backbone RMSD; b) normalised B-factors and the sampling variance on carbon-alpha of each residue. A Pearson correlation > 0.5 was observed between backbone RMSD and sampling variance while no strong correlation between normalised B-factors and variance.

## Footnotes

1 https://github.com/instadeepai/FrameDiPT

2 https://www.rcsb.org/docs/programmatic-access/file-download-services

3 https://github.com/huhlim/cg2all

4 https://www.modelarchive.org/

5 https://biopython.org/

6 TCR:pMHC class I structures missing MHC beta chains were kept, since the domain is not involved in the TCR:pMHC interface.

